# Tracking strains predicts personal microbiomes and reveals recent adaptive evolution

**DOI:** 10.1101/2020.09.14.296970

**Authors:** Shijie Zhao, Chengzhen L. Dai, Ethan D. Evans, Ziyu Lu, Eric J. Alm

## Abstract

An individual’s microbiome consists of a diverse set of bacterial strains that encode rich information on its colonization and evolutionary history. Here, we introduce a versatile and straightforward reference-based strain tracking approach (StrainTrack) that determines whether distinct metagenomes carry closely-related strains based on gene presence and absence profiles. We show that StrainTrack can predict whether two metagenomes originate from the same donor via counting the number of species sharing closely-related strains, achieving >96% specificity, and ∼100% sensitivity. When applied to the metagenomes of adult twins in the TwinsUK registry, we identify six cases of closely-related strains carried by both twins, potentially over decades of colonization.

## Introduction

The human gut microbiome harbors a complex community of microbial species stably colonizing for years or even decades (Faith *et al*., 2013; Lloyd-Price *et al*., 2017). While individuals from the same human population usually contain similar species, different people typically carry person-specific strains, distinguished by genomic variations such as single nucleotide polymorphisms (SNPs), insertion/deletions, and gene presence/absence (Scholz *et al*., 2016; Truong *et al*., 2017). In principle, tracking closely-related strains paves the way to understanding how microbial strains are transmitted between family members, across social networks, after fecal microbiota transplantation (FMT), and throughout an infection (Ferretti *et al*., 2018; Smillie *et al*., 2018; Brito *et al*., 2019). Such fine-scale monitoring of strains can provide more significant insights into how the microbiome responds to host and environmental factors, with potential applications in the personalization of treatment such as FMT donor selection (Duvallet *et al*., 2017).

Various computational methods are available for strain-level analysis of metagenomic samples. One class of methods resolve strains via identifying SNPs between different metagenomes by aligning short reads to species reference genomes (Luo *et al*., 2015; Costea *et al*., 2017; Truong *et al*., 2017; Smillie *et al*., 2018). However, calling SNPs is prone to false positives, as high-resolution strain profiling usually requires careful and iterative tuning of the filtering parameters (Brito and Alm, 2016). In addition, when multiple strains coexist, it is difficult to phase alleles carried by different strains. Another commonly-used approach involves the analysis of pangenomes, particularly the accessory genomic regions carried by distinct strains. Unlike SNP-based methods, pangenome-based approaches are more robust to parameter changes, yet they require an extensive database of pangenomes built from a diverse set of available strains and substantial computational resources; to date, only a small number of gut species have high-quality pangenomes available (Zhu *et al*., 2015; Scholz *et al*., 2016). Overall, these approaches are limited in their ability to leverage the strain-level information to compare the overall similarity of metagenomes samples and identify personalized signatures, as they often only distinguish strains from one bacterial species at a time. A previous attempt at differentiating microbiome samples and identifying personalization using metagenomic codes was only able achieve 80% accuracy (Franzosa *et al*., 2015).

Here, we introduce a flexible and simple method—StrainTrack—that uses a single reference genome for individual species to compare strain identities between metagenomes. We assume that the accessory genomes of microbial species are highly individual-specific and that many strains can stably colonize a microbiome for years (Faith *et al*., 2013; Zhao *et al*., 2019). We compare the accessory genomes of 40 prevalent gut bacteria species using public metagenomes and demonstrate evidence supporting this assumption for 25 out of the 40 species. We further design a classification rule that leverages these comparisons to predict whether two metagenomes belong to the same individual. We achieve near-perfect specificity and sensitivity when applied to the metagenomes of the Human Microbiome Project (HMP) and the Broad Next 10 (BN10) project. In doing so, we find evidence supporting a mislabeling of donor IDs in a pair of the HMP metagenomes. Lastly, when StrainTrack is applied to track strains from adult twins, we find that certain twins can share strains that may have colonized for potentially decades and discover signatures for adaptive evolution in these shared strains.

## Results

### Accessory genome difference (AGD) as a metric to define inter-sample strain variance

We develop a bioinformatic workflow (StrainTrack) to achieve strain-level comparisons between metagenomic samples by aligning short reads against a set of well-assembled reference genomes. In particular, we align metagenomic reads against a curated collection of 40 abundant or well-studied representative gut bacterial species (Lloyd-Price et al., 2017; Xie et al., 2016, **Table S1**). To quantify the inter-sample strain difference, we develop a metric that estimates the fraction of the reference genome that is variable between the two metagenomes. For any given species, we calculate the relative sequencing depth for every 5kb genomic window. We then compare the relative sequencing depth of each genomic window between sample pairs (Methods). If the average sequencing depth of a genomic window is >50% of the average depth in one sample and with <5% of the average depth in another sample, we designate this genomic region as a differential region. We define the fraction of the reference genome that is designated as differential regions as the accessory genome difference (AGD; Methods).

To demonstrate how AGDs can reveal personalized strain signatures, we examine *Bacteroides vulgatus*, a prevalent species that inhabits the large intestine (Yatsunenko *et al*., 2012). As an example, a pair of distinct metagenomic samples from the same HMP donor has an AGD of 0 (**Figure 1A, 1B**), while a pair of metagenomes from two distinct subjects shows an AGD of 0.040 (**Figure 1C**). We estimate AGDs for all pairwise HMP metagenomes for *B. vulgatus* and observe a clear difference between the inter-subject and intra-subject AGD profiles (Methods; **Figure 1D**). Further supporting this sharp difference, we generate a receiver operating characteristic curve and calculate the area under the curve (AUC) to be 0.989 (**Figure 1E**). We pick an AGD cutoff that maximizes the sum of sensitivity and specificity (Methods) for *B. vulgatus*. StrainTrack then determines that two metagenomes have a personalized signature for *B. vulgatus* if the inter-sample AGD is smaller than this cutoff. We expanded this AGD comparison to all 40 species, obtaining an AUC and a species-specific AGD cutoff for each species (**Figure S1, Table S1**). Among these 40 species, 25 of them have an AUC higher than 0.975.

**Figure 1.**
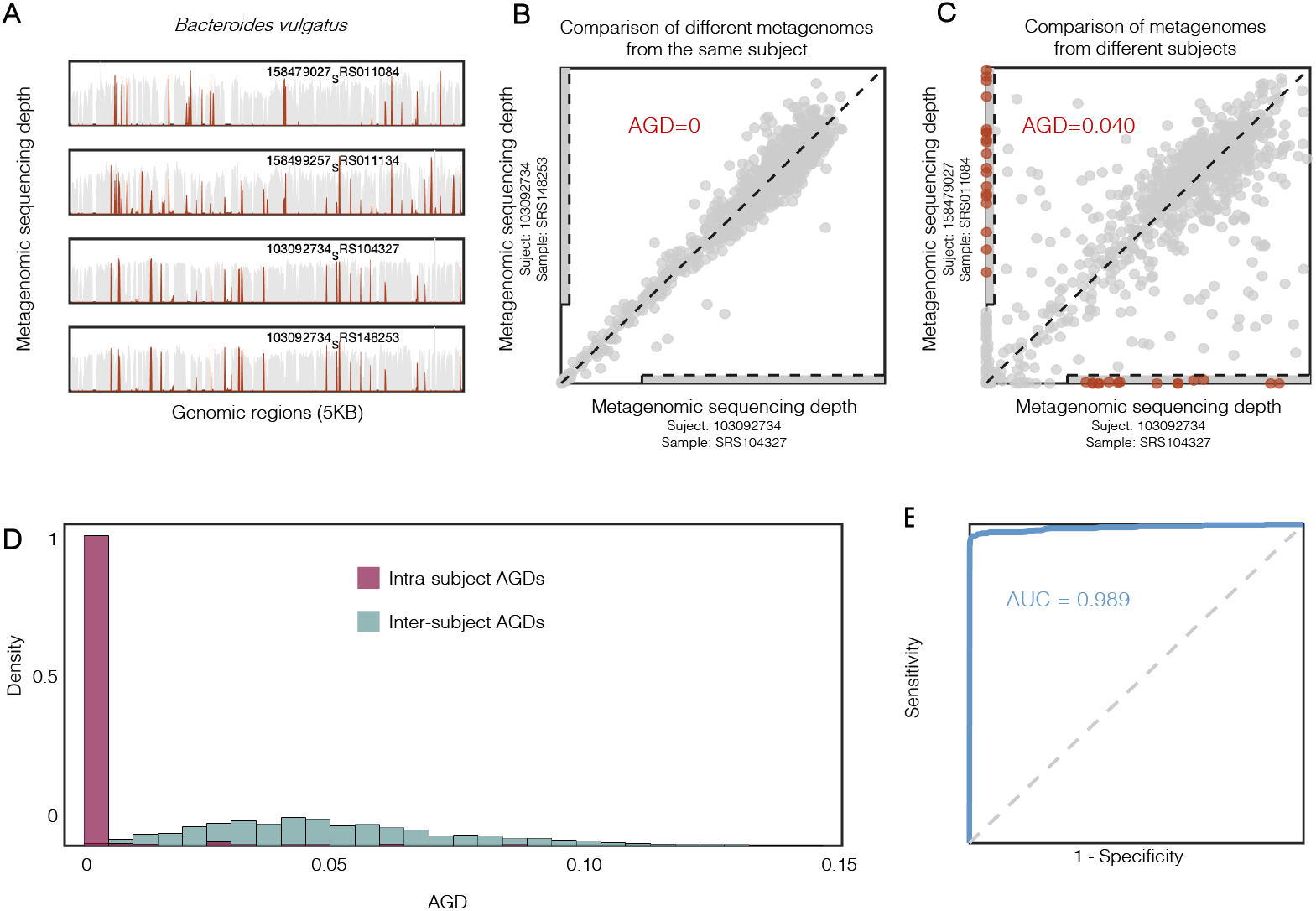
Accessory genome difference (AGD) as a metric to define inter-sample strain variance for *B. vulgatus*. (**A**) Examples showing that *B. vulgatus* strains from distinct subjects differ in accessory genomes (top three panels) and strains from the same subject share similar accessory genomes (bottom two panels). Sequencing depths over the *B. vulgatus* reference are presented for four HMP metagenomes. Genomic regions that are differentially present between the samples are colored in red; genomic regions that are present in all four metagenomes are colored in gray. (**B**) Graphical illustration of calculating AGD for *B. vulgatus* for a pair of metagenomic samples from the same subject. Each dot represents the sequencing depth of a 5 Kb genomic window. (**C**) Graphical illustration of calculating AGD for *B. vulgatus* for a pair of metagenomic samples from two different subjects. Each dot represents the sequencing depth of a 5 Kb genomic window. Genomic regions that are differentially present between the two samples are colored in red. (**D**) Density histograms for intra-subject AGD profile (red) and inter-subject AGD profile (green) of *B. vulgatus*. (**E**) ROC analysis for the AGD profiles of *B. vulgatus*. To obtain sets of sensitivity and specificity to draw the curve, we set cutoffs from 0 to 1 with 0.0001 intervals.

### StrainTrack predicts personal microbiomes for distinct people

Previous reports show that a pair of metagenomes from the same individual share strains for multiple species, while a pair of unrelated metagenomes are unlikely to harbor closely-related strains (Franzosa *et al*., 2015; Lloyd-Price *et al*., 2017). We, therefore, reason that StrainTrack results could be harnessed to predict whether different metagenomic samples are from the same donor based on the number of species sharing personalized genomic signatures. To do so, we design a classification rule that labels two metagenomic samples are from the same donor if more than two species share personalized signatures (**Figure 2**, Methods). To minimize false predictions for individual species, we only considered the 25 species that displayed AUC values higher than 0.975 in the AGD analysis (**Figure S1**; **Table S1**).

**Figure 2.**
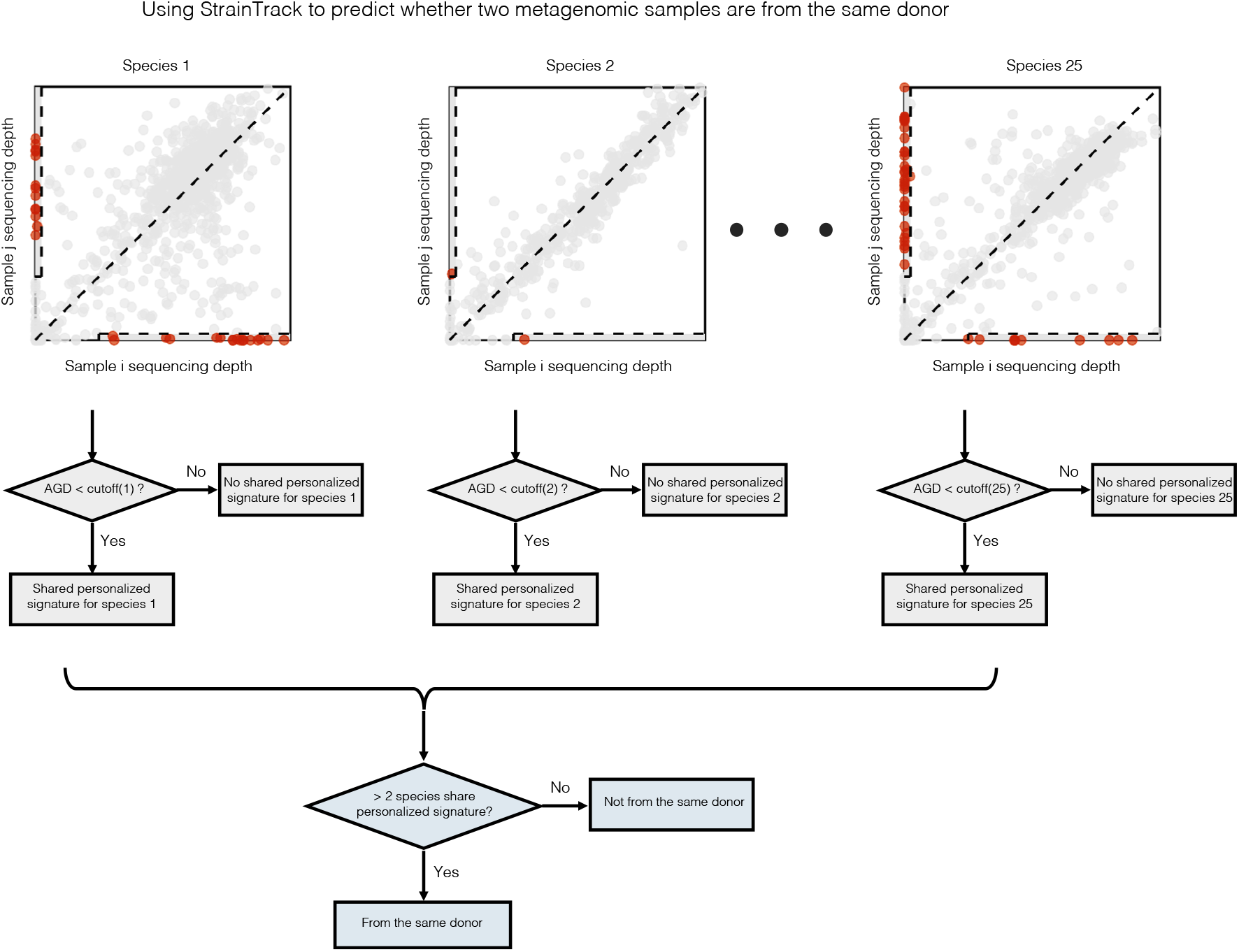
StrainTrack predicts personal microbiomes for distinct people. For a given pair of metagenomes, an inter-sample AGD is calculated for each of the 25 species that show substantial differences between intra-subject and inter-subject AGD profiles (**Table S1**). For each species, the inter-sample AGD is compared to the species-specific AGD cutoff (**Table S1**) and StrainTrack predicts that the metagenomes share the personalized signature for this species if the AGD is smaller than the species-specific cutoff. When more than two species share personalized signature between the two metagenomes, StrainTrack predicts that these two samples belong to the same stool donor.

To examine the performance of this StrainTrack-based classifier, we use it to predict donors for all pairs of 535 HMP metagenomic samples. These samples are from 250 distinct subjects, of which 161 of them had more than one sample (Lloyd-Price *et al*., 2017). When comparing all pairwise samples from HMP, our classifier provides us with a sensitivity of 95.79% and a specificity of 99.99% (**Figure 3A**, Methods). To validate this classifier with an independent test dataset, we apply StrainTrack to the Broad Next 10 dataset (Poyet *et al*., 2019), consisting of 410 metagenomic samples from 50 distinct individuals, and achieve a 100% specificity and 100% sensitivity (**Figure 3B**).

**Figure 3.**
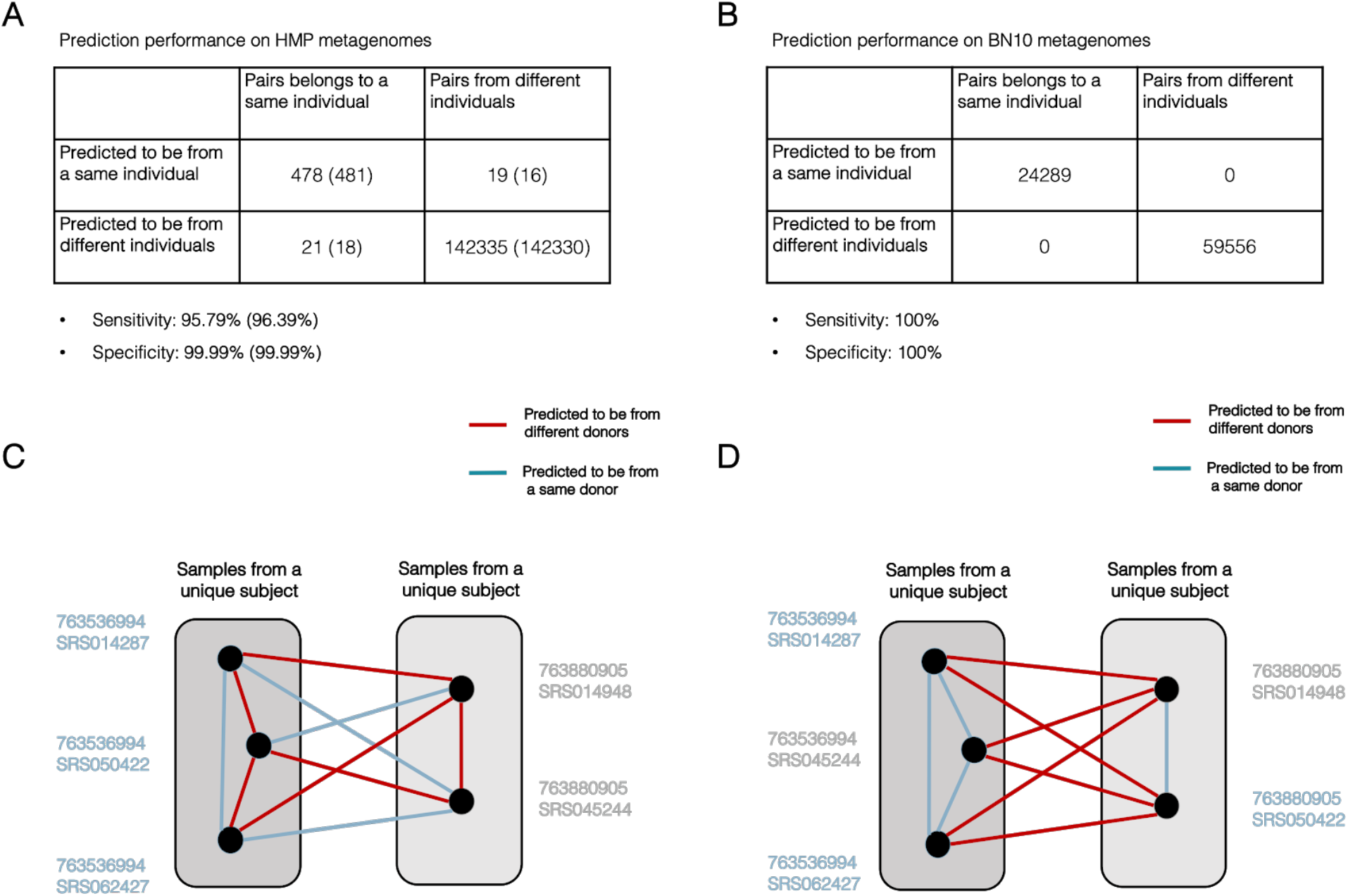
StrainTrack-based classifier achieves >96% sensitivity and ∼100% specificity in predicting metagenome donor. (**A**) Contingency table shows the results of using StrainTrack to predict if a pair of HMP metagenomes belong to the same donor. Numbers in parentheses are after correcting for misclassifications due to the putative mislabeling illustrated in (**C**) and (**D**). (**B**) Contingency table shows the results of using StrainTrack to predict if a pair of BN10 metagenomes are from the same donor. (**C**) Predictions for a set of five HMP metagenomes with proposed misclassifications. Samples labeled by HMP from the same donor are grouped into the same block. Given the empirical false negative rate (< 5 %) and false positive rate (< 0.02 %), the probability of observing these predictions is ∼10^−16^ (false positive rate^3^ * true positive rate^1^ * false negative rate^3^ * true negative rate^3^) (**D**) If the donor IDs of sample SRS050422 and SRS045244 are flipped, the predicted results are consistent with expectations.

From the HMP dataset, we notice a potential mislabeling of donor IDs for a pair of metagenomic samples. We observe that the sample SRS045244 from subject 763880905 is predicted by StrainTrack to share a microbiome donor with two samples from subject 763536994 (SRS014287 and SRS062427); meanwhile, a different sample from the same subject (763880905, sample SRS014948) is matched to the remaining sample from subject 763536994 (SRS050422, **Figure 3C**). Given that the empirical estimation of false negative rate is < 5% and false positive rate is < 0.02% (**Figure 3A**), we estimate that the probability of observing these donor matching patterns is <10^−16^ (**Figure 3C**). However, if we assume that the donor labels of sample SRS050422 and SRS045244 were shuffled, the prediction results are identical to expectations (**Figure 3D**). This analysis suggests that there is a mislabeling in the HMP metagenomes and we offer a parsimonious solution to correct it. After fixing this putative label-shuffling, StrainTrack-based classifier displays an updated sensitivity of 96.4% for the HMP metagenomes.

### Family members, especially young children, share many closely-related strains

Both HMP and BN10 donors consist of mostly unrelated individuals from the US, and it is therefore expected that distinct donors carry distinct strains. However, members from the same household may share a substantial level of strains, especially for children, whose microbiome may partly derive from their parents (Ferretti *et al*., 2018). To test whether StrainTrack can be applied to people from the same family, we examine the metagenomes of an 8-member family, consisting of a mother, father, and six children of ages 0, 2, 4, 6, 8, and 10 years old (Schloss *et al*., 2014). Each family member has between 1 and 3 metagenomic samples sequenced (**Figure S2**). While most pairs of family members can be successfully separated by StrainTrack, we find that certain family members shared closely-related strains that complicate the classification accuracy. In particular, the microbiomes of the 4, 6 and 8-year old children were predicted to be from the same individual (**Figure S2**). These three children shared similar strain signatures for multiple species, including many *Bifidobacterium* and *Bacteroides* strains. While such results suggest that our method is limited in differentiating the microbiomes of family members, they demonstrate the ability of StrainTrack in identifying transmission events between subjects (Brito *et al*., 2019).

### Adult twins share closely-related strains at low frequency, potentially for decades

We next explore the ability of StrainTrack to detect strains shared by different human subjects over a longer period of time (with potential transmission events). We select a dataset from adult twins in the UK Twin Registry, including 125 pairs of adult twins between 50 and 70 years old (Xie *et al*., 2016). We first calculated the inter-twin AGDs for the 25 species that are used in the StrainTrack predictor. While the majority of the species do not have inter-twin AGDs smaller than the species-specific cutoffs, we identify 27 cases in which a species shared a personalized signature between twins (**Table S2**).

To validate if these identified personalized signatures reveal real transmission events between the twins, we examine the evolutionary history of these strains to rule out apparent false positives. We identify the genome-wide distribution of SNPs for these strains by searching for nucleotide positions in which the major alleles are discordant between twins (Methods). We exclude species with a signature of multiple-strain colonization, defined by an excess of genomic positions with major allele frequency smaller than 0.95, were excluded (Methods). We also excluded twin-species combinations containing genomic regions with more than 20 SNPs/Kb (Methods, **Table S2, Figure S3**). Given that the documented molecular clock for bacterial species in natural environments range from 0.5–5 SNPs/year (Didelot *et al*., 2016), these numbers of SNPs are inconsistent with recent transmission events and are likely distinct colonization events by closely-related strains or transmission complicated by homologous recombination. After filtering, there were six cases of shared, closely-related strains showing evidence of recently-emerged mutations. Our analysis suggested that between 4 and 74 SNPs separated the strains harbored by distinct twins (**Figure 4A-B, Table S2**). It is worth noting that our SNP analysis can only detect mutations that reached high frequency within either twin’s microbiome. Given the molecular clocks for bacterial species, these levels of SNPs suggest years to decades of evolutionary divergence between the twins. Since these twins have mostly been living apart for 30–50 years, it is thus likely that these mutations emerged and accumulated independently within the gut of each twin, while some shared strains may have been colonizing both twins for decades.

**Figure 4.**
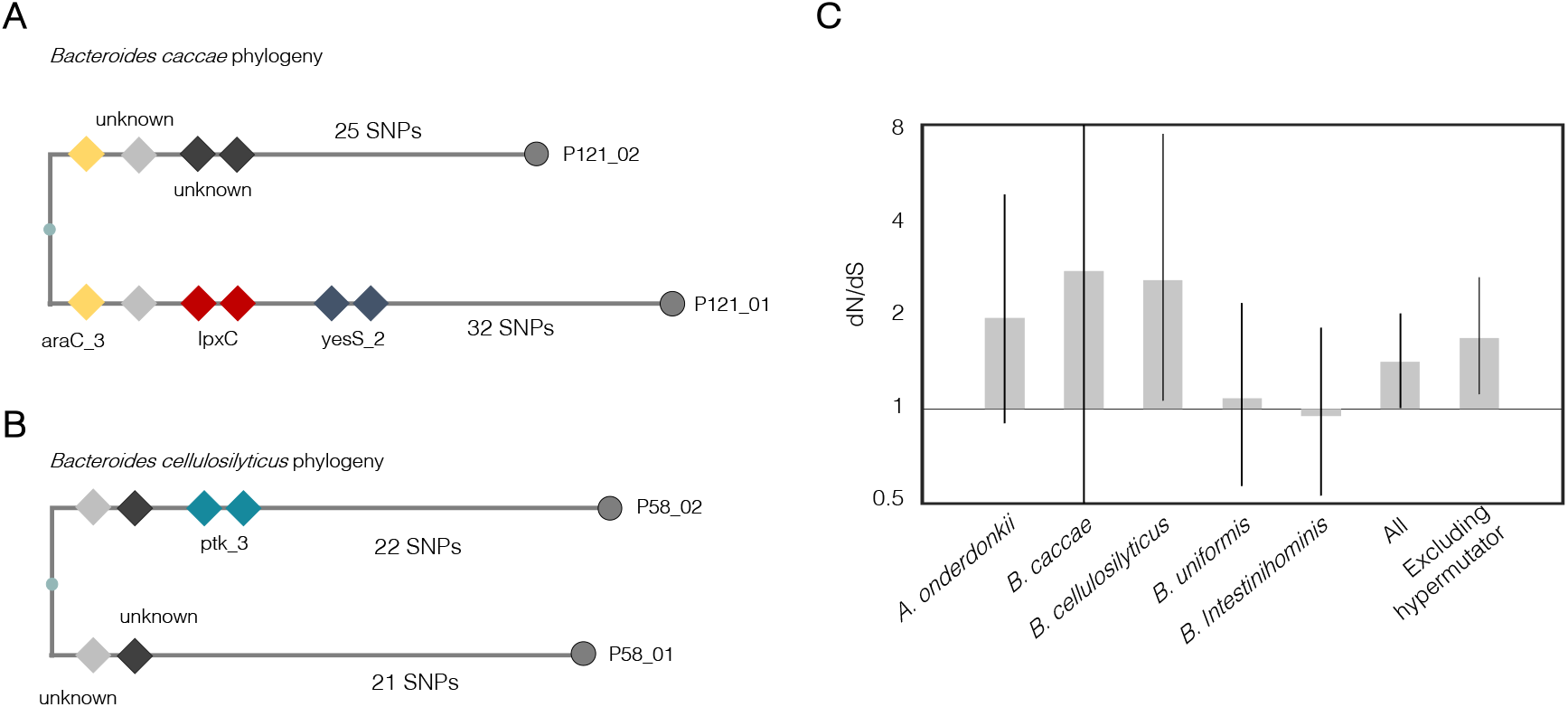
Strains shared by twin pairs show signatures of adaptive within-person evolution. (**A**) and (**B**) Phylogenies of a *B. caccae* strain shared between the twin pair P121 and a *B. cellulosilyticus* strain shared between the twin pair P58. Diamonds represent mutations in genes with more than one SNP identified between the twins. Distinct genes are colored differently and annotated. (**C**) dN/dS is calculated for SNPs identified in each individual species and combined. dN/dS is also calculated for combined SNPs excluding *B. intestinihominis*, as this species appears to be a hypermutator. Error bars represent the 95% confidence intervals.

### Strains shared by twin pairs show a signature of adaptive within-person evolution

The shared strains between adult twins and the recently emerged SNPs (years to decades) provided an opportunity to investigate the within-person evolutionary process of these strains. To examine if these point mutations reflect adaptive evolution within the twins, we calculated dN/dS for the six strains that showed evidence for recent transmission (these six strains are from five species). dN/dS is the normalized ratio of non-synonymous mutations to synonymous mutations and is a canonical measure of selection (Methods). For the five species tested, we found the values dN/dS are larger than or very close to one (**Figure 4B**). When combining SNPs identified from all five species, we obtained a dN/dS that is significantly bigger than one, suggesting genome-wide adaptive evolution dominates the within-person evolution for these species. We thus suggest that the mutations that swept in these twins were driven by adaptive evolution.

## Discussion

Here, we introduce a new metagenomic analysis framework (StrainTrack) for rapid identification of closely-related strains between metagenomes and accurate classification of shared microbiomes. Compared to other strain-tracking methods, StrainTrack has a simple implementation and is easy to be modified for different tasks. Leveraging the assumption that unrelated humans carry strains with unique accessory genome profiles, we built a StrainTrack-based classifier with 25 species that have distinct intra-personal and inter-personal AGD profiles. Such differences allow us to infer with confidence whether two metagenomic samples share strains and further predict personal microbiomes with high accuracy. StrainTrack performs particularly well for HMP and BN10 metagenomes, with only a few cases of misclassification in HMP samples. Our sensitivity and specificity are better compared to a previously reported classifier with 80% accuracy in recovering HMP microbiome donors (Franzosa *et al*., 2015). In addition, we find one case that the donor IDs of a pair of metagenomes appeared to have been mislabeled. This mislabeling has been hinted in the supplementary materials from a previous report (Schloissnig *et al*., 2013).

Our method also enabled us to track closely-related strains across metagenome samples and helped identify strains shared by twin pairs, potentially over decades of colonization. Further analysis of the point mutations between the twin pairs revealed evidence that these shared strains experience genome-wide adaptive evolution. Our analysis only accounted for mutations that nearly sweep either twin and is likely missing mutations that are present at medium or low frequencies. In addition, we identified six shared strains all from the *Bacteroidetes* phylum. Nonetheless, our results demonstrated that adaptive evolution might dominate at short timescale across the genome. This is striking given compelling evidence that purifying selection dominates evolution at timescales of tens of thousands of years (Schloissnig *et al*., 2013; Garud *et al*., 2017; Zhao *et al*., 2019). To solve this discrepancy, we propose two theoretical scenarios to reconcile signals from the two timescales. One possibility is that many strains carried by an individual will be lost over transmission between human populations; thus, within-person adaptive mutations are rarely transmitted to new human hosts. Another possibility is that within-person adaptive mutations are person-specific and usually lead to selective disadvantages in new human hosts, and over time, these adaptive mutations will be selected against by natural forces. Future studies with larger sample sizes and more strains from diverse taxonomic groups are needed to test these hypotheses.

## Acknowledgements

We thank Fangqiong Ling for inspiring us to explore predicting human host using metagenomic data. We appreciate Sean Gibbons for insightful discussions throughout the development of this paper. We are grateful to Tami Lieberman, Michael Baym, Lei Dai, Sean Kearney, Thomas Gurry, and Xiaoqian Yu for the helpful and constructive discussions. We are thankful to Xiaofang Jiang, Mathieu Groussin and Mahilde Poyet for assistance in obtaining and preprocessing BN10 metagenomes.

## Declaration

### Author contributions

S.Z. and E.J.A design the study; S.Z. leads the metagenomic analysis with contributions from C.L.D, Z.L., W.Z., A.Z. and E.D.E.; S.Z. writes the manuscript with significant inputs from C.L.D, E.D.E., and E.J.A.

## Methods

### Metagenomic datasets used in this study

We consider three publicly available datasets for this study: the Human Microbiome Project (Lloyd-Price *et al*., 2017) (535 samples from 250 subjects: https://www.hmpdacc.org), the TwinsUK study (Xie *et al*., 2016) (250 samples from 250 subjects; ERP010708), and a gut microbiome study of an eight-member family (Schloss *et al*., 2014) (15 samples from 8 family members). We also include datasets from the Broad Next 10 project (410 samples from 50 subjects; PRJNA544527) (Poyet *et al*., 2019).

### Reference genomes

The accessory genome comparison requires that a strain has adequate sequencing depths from both metagenomic samples. To meet this criterion, we manually curate species that are abundant and prevalent in the HMP and TwinsUK datasets and include some well-characterized species found in the gut microbiome (e.g., *E. coli*), totalling 40 species. A representative reference genome for each species was obtained from NCBI, and a single fasta file was generated that contains these 40 genomes. To simplify the downstream analysis, for references with multiple scaffolds, we connect the sequences from different scaffolds to form a single artificial contig. The list of references used in this study can be found in **Table S1**.

### Metagenomic reads alignment

Metagenomic reads are trimmed and filtered using Cutadapt and Sickle Sickle (Joshi and Fass, 2011; Martin, 2011). The filtered reads are aligned to the combined reference genome using Bowtie2 (Langmead and Salzberg, 2012) (Parameters: -X 2000, --no-mixed, --very-sensitive, -- n-ceil 0,0.01, --un-conc). Alignment files (sam format) are converted to pileup files using Samtools (Li *et al*., 2009) (Step 1: samtools view -bS; Step 2: samtools sort; Step 3: samtools mpileup -q30 -x -s -O -d3000). From these pileup files, we extract information about read depth at each genomic position from the combined reference.

### AGD calculation

AGD is defined as the fraction of accessory genomic regions that are different between two metagenomic samples. AGD is used to quantify the strain-strain distance between a pair of metagenomes. We first divide the single contig (see Methods: Reference genomes) for the targeted species into 5 Kb genomic windows. Average sequencing depth was calculated for each of the genomic windows from both metagenomes. A genomic region is designated as different when its sequencing depth is lower than 5% of the average sequencing depth in one sample and is higher than 50% of the average in the other sample. To avoid inaccurate estimation of average sequencing depth, due to abnormal alignment at mobile genomic regions and regions in the reference genome that are not present in the sample-specific strains, average sequencing depth is defined as the mean depth of genomic positions with sequence depth between the 25 and 75 percentiles of genome-wide sequencing depth.

For each species, we generate a cutoff that maximizes the differentiation of inter-subject and intra-subject AGD profiles (**Figure 1D, Table S1**). This is accomplished by finding a cutoff that maximizes the sum of sensitivity and specificity.

### StrainTrack-based classifier

StrainTrack predictor includes 25 species that the AGD analysis shows AUC > 0.975. These species all have sharply distinct intra-subject and inter-subject AGD profiles that allow detecting closely-related strains with low degrees of false prediction. For a given pair of metagenomes, we perform metagenomic alignment to the 25 species references and calculat the AGD for each species between the two metagenomes. For each of the 25 species, the AGD is compared to the species-specific AGD cutoff (**Table S1**). If AGD is smaller than the cutoff, the species is classified as having a personalized signature between the two samples. If more than two species share a personalized signature between the two metagenomes, StrainTrack predicts that the two samples belong to the same stool donor (**Figure 2**).

### Identification of mutations between twins

For each species that share personalized signatures between a twin pair, candidate SNPs are identified using SAMtools and filtered using filters optimized from previous work (Lieberman *et al*., 2011, 2014). In particular, genomic positions were considered to be potential SNP positions if the twins were discordant on the called base and both samples had: FQ score less than 30, at least 1 read aligning either forward strand or reverse strand and a major allele frequency of at least 80%. The median coverage across samples must be more than one read. Samples with potential multiple-strain colonization are discarded in the analysis (>3% of the variable positions have <95% major allele frequency, [reference]). In addition, regions that are not within 50-200% of the average sequencing depth of the genome are discarded, as these polymorphisms are likely from species that share homologous sequence to the reference. Detailed information of between-twins SNPs for the shared strains are listed in **Table S2**.

### dN/dS

Mutations were categorized as synonymous (S) or non-synonymous (N) based on open-reading frame annotations from the GenBank files of the reference genomes. To calculate dN/dS for sets of *de novo* mutations (**Figure 4, Table S2**), we normalized the observed N/S ratios by the expected N/S ratios (Zhao *et al*., 2019). For any given set of SNPs, we calculated the expected N/S for these SNPs, accounting for both (1) the different probabilities of acquiring non-synonymous mutations for different types of mutations and (2) the codon compositions of the genes in which these SNPs occurred.

## Supplementary Figures

**Figure S1.**
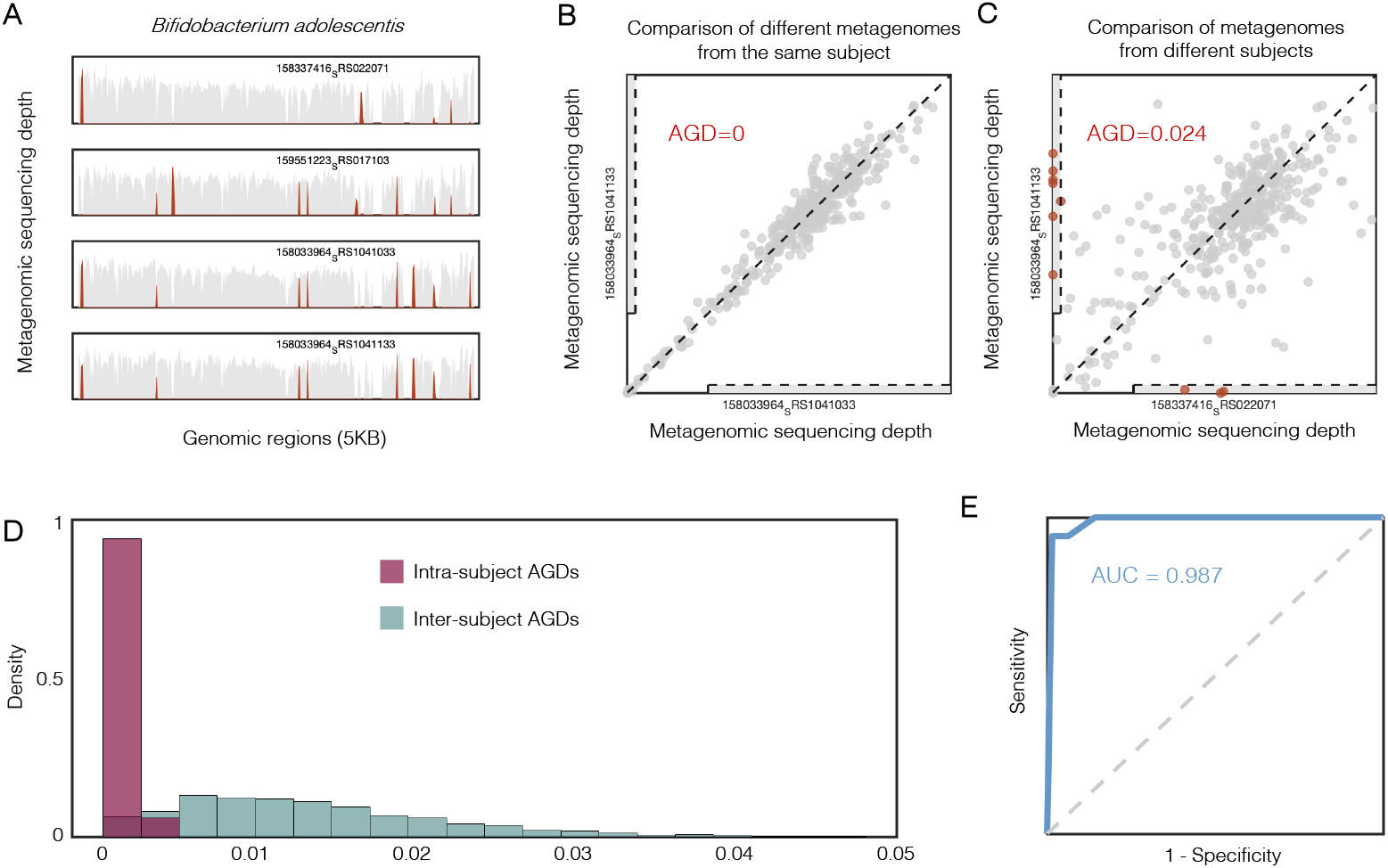
Accessory genome difference (AGD) as a metric to define inter-sample strain variance for *B. adolescentis*. (**A**) Examples showing that *B. adolescentis* strains from different human subjects are different in accessory genomes. Sequencing depths over the *B. adolescentis* reference are presented for four HMP metagenomes. Genomic regions that are differentially present between the samples are colored in red; genomic regions that are present in all four metagenomes are colored in gray. (**B**) Graphical illustration of calculating AGD for *B. adolescentis* for a pair of metagenomic samples from the same subject. Each dot represents the sequencing depth of a 5 Kb genomic window. (**C**) Graphical illustration of calculating AGD for *B. adolescentis* for a pair of metagenomic samples from two different subjects. Each dot represents the sequencing depth of a 5 Kb genomic window. Genomic windows that are differentially present between the two samples are colored in red (Methods) (**D**) Density histograms for intra-subject AGD profile (red) and inter-subject AGD profile (green) of *B. adolescentis*. (**E**) ROC analysis for the AGD profiles of *B. vulgatus*. To obtain (sensitivity, specificity) sets to draw the curve, we set cutoffs from 0 to 1 with 0.0001 intervals.

**Figure S2.**
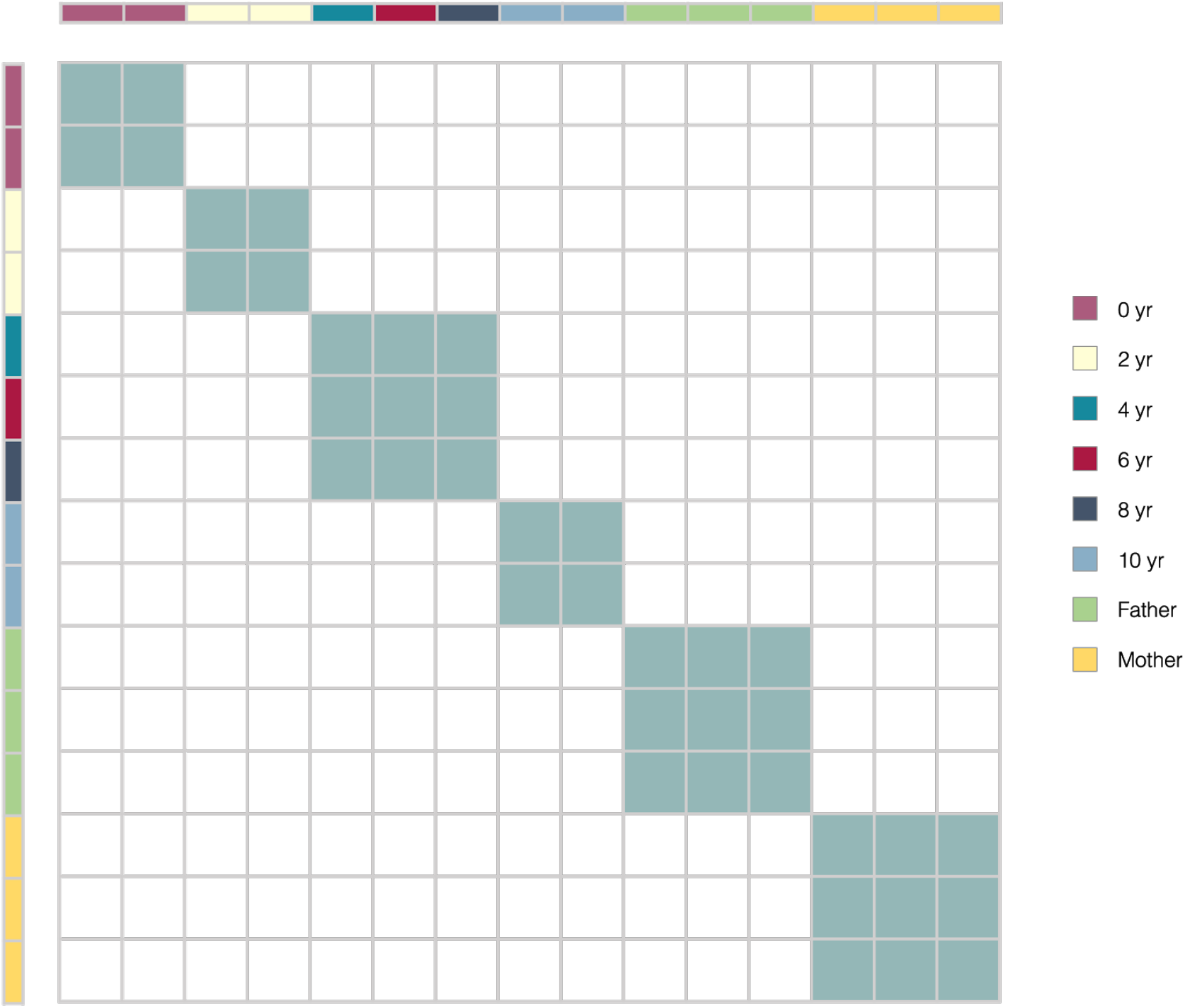
Samples from close family members can be clustered together. StrainTrack is applied to 15 metagenomes from 8 family members. Columns and rows represent distinct metagenomic samples. Row labels and column labels represent the identity of the family member. If two metagenomes are predicted by StrainTrack to be from the same donor, they are colored with green in the heatmap. We notice that StrainTrack cannot distinguish metagenomes from the 4-year old, 6-year old and 8-year old children.

**Figure S3.**
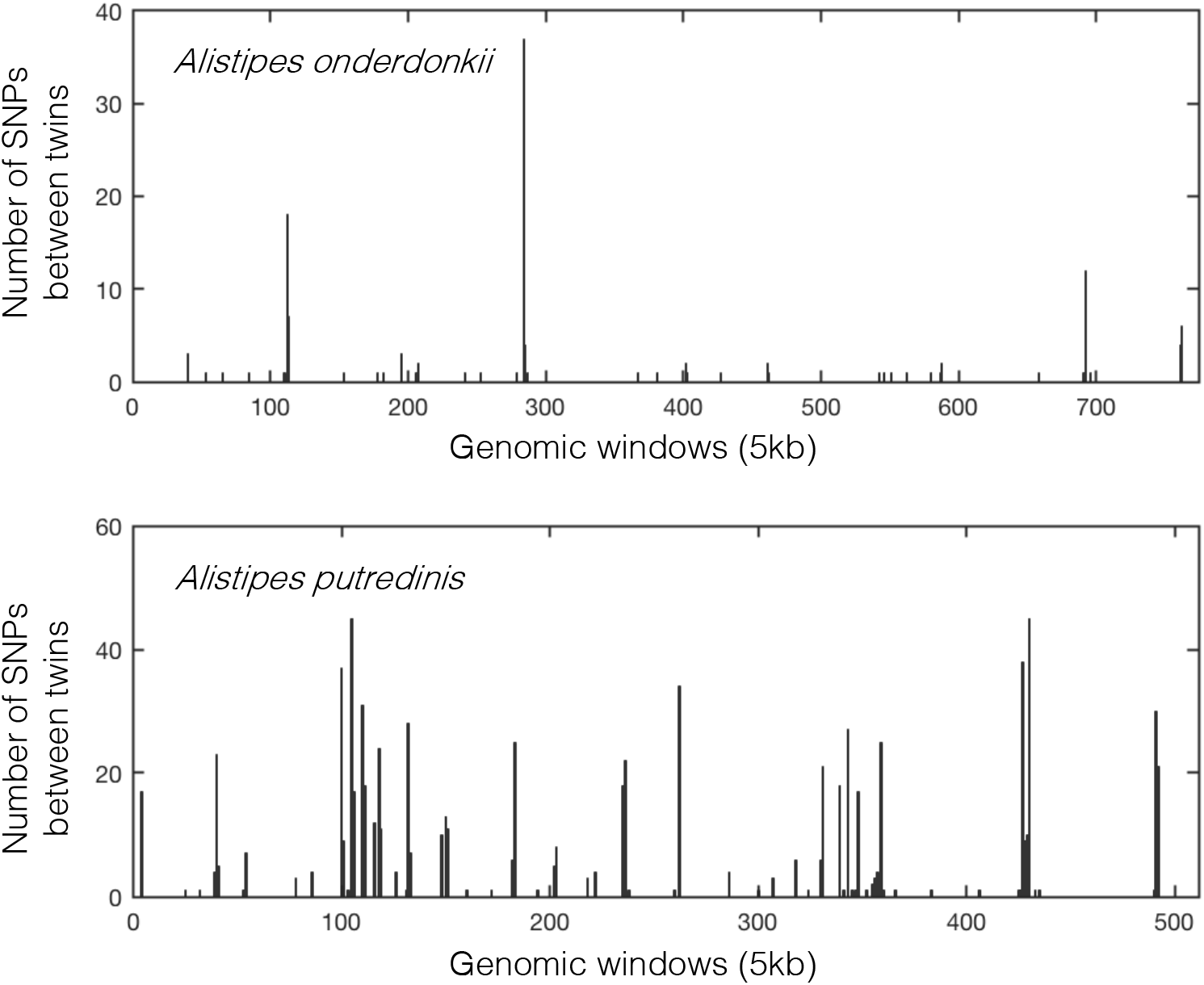
*Alistipes onderdonkii* and *Alistipes putredinis* between UK twin pair P126 show signs for recombination or separate colonization events by closely-related strains. The *Alistipes onderdonkii* and *Alistipes putredinis* strains predicted by StrainTrack as having personalized signature between the twins. Both genomes contain regions enriched for SNPs (>20 SNPs/Kb), suggesting that these two species underwent homologous recombinations or they are not closely-related strains.

**Table S1:**
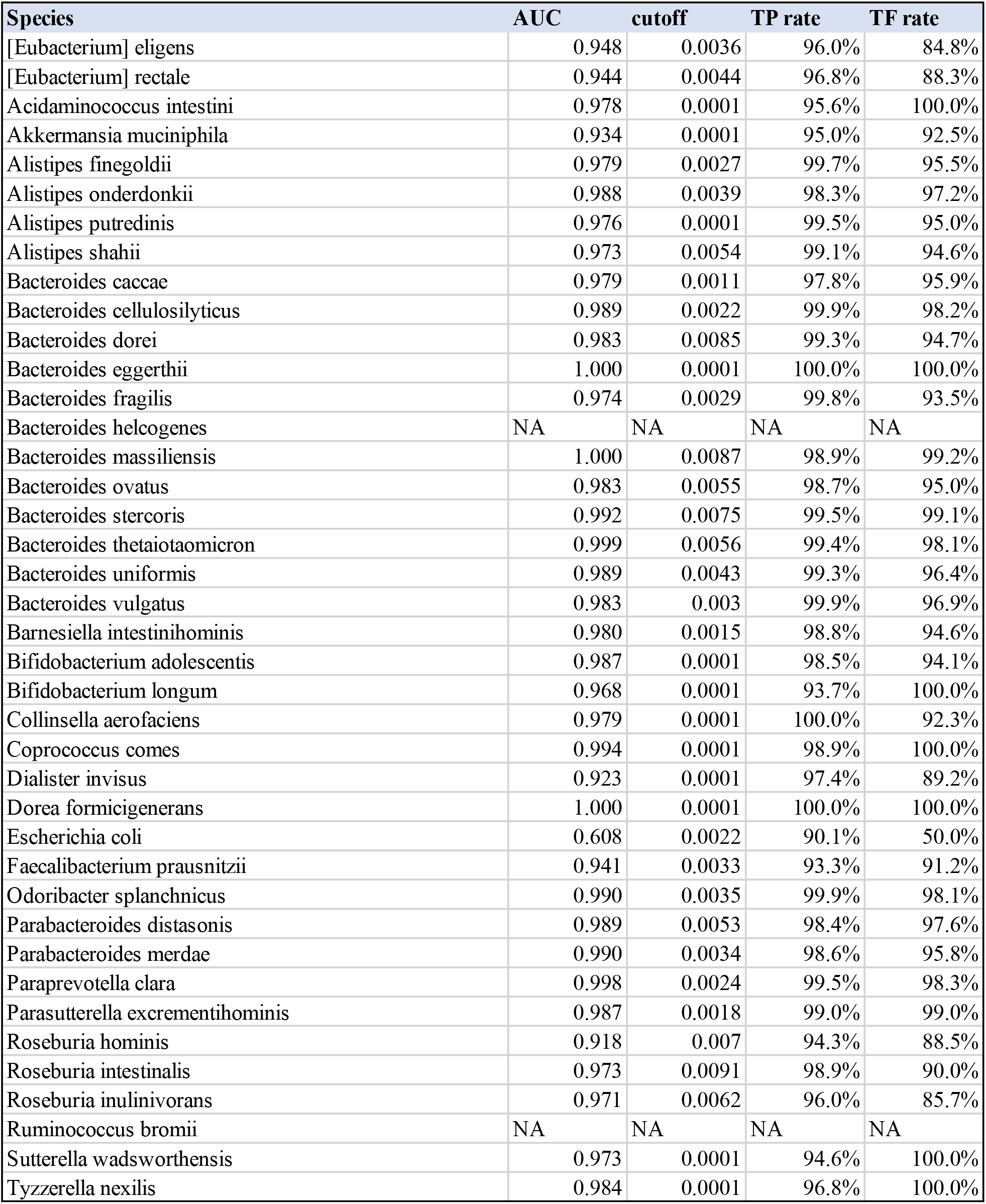
Species used in StrainTrack.

**Table S2:** closely-related strains shared between adult twins.

| **Twin Pairs** | **Species** | **Conclusion from SNP analysis** | **Number of SNPs swept in twin 1** |
| --- | --- | --- | --- |
| P64 | Alistipes onderdonkii | Possibly mixed strains | NA |
| P97 | Alistipes onderdonkii | Closely-related strains | 8 |
| P113 | Alistipes onderdonkii | Possibly mixed strains | NA |
| P119 | Alistipes onderdonkii | Possibly mixed strains | NA |
| P126 | Alistipes onderdonkii | Possibly with recombination | NA |
| P126 | Alistipes putredinis | Possibly with recombination | NA |
| P121 | Bacteroides caccae | Closely-related strains | 57 |
| P58 | Bacteroides cellulosilyticus | Closely-related strains | 43 |
| P73 | Bacteroides ovatus | Possibly mixed strains | NA |
| P15 | Bacteroides uniformis | Closely-related strains | 4 |
| P19 | Bacteroides uniformis | Possibly mixed strains | NA |
| P27 | Bacteroides uniformis | Closely-related strains | 56 |
| P29 | Bacteroides uniformis | Possibly mixed strains | NA |
| P35 | Bacteroides uniformis | Possibly mixed strains | NA |
| P54 | Bacteroides uniformis | Possibly mixed strains | NA |
| P89 | Bacteroides uniformis | Possibly mixed strains | NA |
| P111 | Bacteroides uniformis | Possibly mixed strains | NA |
| P7 | Bacteroides vulgatus | Possibly mixed strains | NA |
| P15 | Bacteroides vulgatus | Possibly mixed strains | NA |
| P29 | Bacteroides vulgatus | Possibly mixed strains | NA |
| P45 | Bacteroides vulgatus | Possibly mixed strains | NA |
| P62 | Bacteroides vulgatus | Possibly mixed strains | NA |
| P70 | Bacteroides vulgatus | Possibly mixed strains | NA |
| P100 | Bacteroides vulgatus | Possibly mixed strains | NA |
| P103 | Bacteroides vulgatus | Possibly mixed strains | NA |
| P15 | Barnesiella intestinihominis | Closely-related strains | 74 |
| P47 | Collinsella aerofaciens | Possibly mixed strains | NA |

